# Regulators of the secretory pathway have distinct inputs into single-celled branching morphogenesis and seamless tube formation in the Drosophila trachea

**DOI:** 10.1101/2021.05.13.443924

**Authors:** Christopher M. Bourne, Daniel C. Lai, Jodi Schottenfeld-Roames

## Abstract

Biological tubes serve as conduits through which gas, nutrients and other important fluids are delivered to tissues. Most biological tubes consist of multiple cells connected by epithelial junctions. Unlike these multicellular tubes, seamless tubes are unicellular and lack junctions. Seamless tubes are present in various organ systems, including the vertebrate vasculature, *C*.*elegans* excretory system, and *Drosophila* tracheal system. The *Drosophila* tracheal system is a network of air-filled tubes that delivers oxygen to all tissues. Specialized cells within the tracheal system, called terminal cells, branch extensively and form seamless tubes. Terminal tracheal tubes are polarized; the lumenal membrane has apical identity whereas the outer membrane exhibits basal characteristics. Although various aspects of membrane trafficking have been implicated in terminal cell morphogenesis, the precise secretory pathway requirements for basal and apical membrane growth have yet to be elucidated. In the present study, we demonstrate that anterograde trafficking, retrograde trafficking and Golgi-to-plasma membrane vesicle fusion are each required for the complex branched architecture of the terminal cell, but their inputs during seamless lumen formation are more varied. The COPII subunit, Sec 31, and ER exit site protein, Sec16, are critical for subcellular tube architecture, whereas the SNARE proteins Syntaxin 5, Syntaxin 1 and Syntaxin15 are required for seamless tube growth and maintenance. These data suggest that distinct components of the secretory pathway have differential contributions to basal and apical membrane growth and maintenance during terminal cell morphogenesis.

## Introduction

Tubulogenesis is a critical process in the formation of a diverse set of organ systems. Biological tubes exist in three forms: multicellular, autocellular, and seamless tubes. Multicellular and autocellular tubes form lumens via cell-cell contacts; unicellular seamless tubes form an intracellular lumen *de novo* in the absence of epithelial cell junctions (Iruela-Arispe and Davis 2009; Lubarsky and Krasnow 2003). Seamless tubes are found in a wide array of organ systems, including the mammalian and zebrafish vasculature, the *C. elegans* excretory system, and the *Drosophila* tracheal system (Sundaram and Cohen 2017). In insects, such as *Drosophila*, oxygen is not carried in circulating hemolymph. Rather a vast network of ramifying tracheal tubes serves to carry oxygen to all tissues (Samakovlis et al. 1996). During network formation, tip cells lead the migration of new branches and align with each other to mediate interconnection of tracheal branches. In this process, tip cells convert themselves into seamless tubes. Likewise, during sprouting angiogenesis in the vertebrate vascular system, endothelial tip cells must ultimately connect new sprouts to other tubes in the vascular network. Seamless tubes are frequently found at the sites of interconnection in the vasculature, suggesting endothelial tip cells also generate seamless tubes (Bär, Güldner, and Wolff 1984). Indeed, the transformation of a tip cell into a “fusion cell” characterized by a unicellular or seamless tube has been shown at these critical junction points in the zebrafish vasculature (Herwig et al. 2011). Since defects in the capillary network have been implicated in a multitude of cardiovascular diseases, and a large percentage of the microvasculature consists of seamless tubes, it is of great interest to uncover the genetic and mechanistic principles underlying seamless tubulogenesis (Sundaram and Cohen 2017; Bär 1980).

Tip cells leading tracheal buds in the fruit fly ultimately differentiate into two specialized cell types that form seamless tubes: fusion cells and terminal cells (Samakovlis et al. 1996). Fusion cells promote anastomosis between branches of the tracheal system; terminal cells located at the termini of tracheal branches ramify extensively on target tissues, like muscle, and act as the principal site of gas exchange (Jarecki, Johnson, and Krasnow 1999). By the end of larval life, terminal cells achieve a complex stellate shape, reminiscent of neurons, but with each cytoplasmic projection having hollowed out to form a tube with apical-basal polarity (Schottenfeld-Roames and Ghabrial 2012; Jones and Metzstein 2011). Membrane addition at the apical domain aids tubulogenesis, whereas growth at the basal membrane promotes branching. The Branchless/Breathless FGF Pathway stimulates the growth of terminal cells and is critical for intracellular lumen formation (Lee et al. 1996; Sutherland, Samakovlis, and Krasnow 1996). Forward genetic screens have revealed a number of factors acting downstream of the FGF signaling pathway in generating this unicellular branched tubular architecture (Ghabrial, Levi, and Krasnow 2011; Jones and Metzstein 2011; Levi, Ghabrial, and Krasnow 2006; Baer, Bilstein, and Leptin 2007). The genes identified thus far have highlighted the importance of apical-basal polarity, the actin and microtubule cytoskeletons, and membrane trafficking during terminal cell morphogenesis, yet the origins of the apical and basal domains remain unclear (Schottenfeld-Roames, Rosa, and Ghabrial 2014; JayaNandanan, Mathew, and Leptin 2014; Lubarsky and Krasnow 2003; Levi, Ghabrial, and Krasnow 2006; Jones and Metzstein 2011; Gervais and Casanova 2010).

*In vitro* cell culture of human endothelial cells has suggested that the lumenal membrane of seamless tubes is endocytic in origin. HUVEC cells plated on a 3-dimensional matrix form an intracellular lumen through the fusion and coalescence of pinocytic vacuoles (Davis and Bayless 2003). Live imaging of seamless tubulogenesis in the intersegmental vessels (ISVs) of the Zebrafish vascular system also supports a role for intracellular vacuoles driving lumen initiation, although the nature of these vacuoles has yet to be elucidated (Yu et al. 2015). In the *C. elegans* excretory canal cell, intracellular tube formation is driven by the alignment of small vacuoles along microfilament tracks, however the targeted addition of membrane from peri-apical canalicular reservoirs is responsible for seamless tube growth (Kolotuev et al. 2013; Khan et al. 2013). Tube formation and maturation in *Drosophila* terminal cells requires the coordinated activity of the microtubule and actin cytoskeleton and vesicle trafficking. Dynein-dependent transport of cargo towards the minus-ends of microtubules is critical for lumenal membrane expansion into branch extensions (Schottenfeld-Roames and Ghabrial 2012; Gervais and Casanova 2010). The identity of this cargo, and whether vesicles from specific membrane compartments in the cell associate with the dynein-motor complex to promote tube extension is still an active area of investigation. Recycling endosomes are one population of vesicles that contribute to lumenal membrane growth. Rab35, an effector of recycling endocytosis, directs the central-to-peripheral polarized growth of the tube, likely via a microtubule transport mechanism (Schottenfeld-Roames and Ghabrial 2012). Syntaxin 7-dependent early endosomal fusion events maintain tube morphology through the regulation of the actin-bundling protein Moesin and the apical actin meshwork (Schottenfeld-Roames, Rosa, and Ghabrial 2014; Hong 2005). The integrity of the actin cytoskeleton at the apical membrane of terminal cells is also mediated through the synaptotagmin-like protein Bitesize, who partners with Moesin to direct the proper deposition of actin and apical determinants at the lumenal, but not basal, membrane of the cell (JayaNandanan, Mathew, and Leptin 2014). Recently, multivesicular bodies have been implicated in nascent tube growth at terminal cell branch points and for membrane allocation to both the apical and basal domains (Nikolova and Metzstein 2015; Mathew et al. 2020).

Although much remains to be elucidated about the trafficking events directing *de novo* lumen formation, even less is understood about the contributions of distinct membrane compartments during terminal cell elongation and ramification. During embryonic development, Sec6 and Sec8 localize to the cell periphery of the elongating terminal cell, with preferential accumulation at the cell tip (Gervais and Casanova 2010). Accordingly, compromising exocyst complex activity leads to a significant reduction in cell size and branch complexity during later larval stages (Jones et al. 2014). Endocytosis plays an equally critical role in sculpting the branched architecture of this cell, as Syntaxin7, Rab5 and Dynamin mutant larval terminal cells are severely pruned (Schottenfeld-Roames, Rosa, and Ghabrial 2014). Polarized trafficking of intracellular compartments has also been shown to be important in neuronal dendrite formation, which is thought to be mediated through specialized golgi outposts at branch points as well as the transport of Rab5-containing early endosomes to distal branches (Ye et al. 2007; Horton et al. 2005; Satoh et al. 2008). These findings collectively speak to the complex crosstalk occurring between membrane compartments as a terminal cell hollows out each branch extension; the importance of understanding the breadth of membrane contributions during such a dramatic cell shape transformation cannot be understated.

To identify target membrane compartments driving seamless tube formation and branching morphogenesis, we turned our attention to the large family of SNARE proteins responsible for mediating membrane fusion during all vesicle trafficking events. We surveyed all Qa and select Qc SNARE vesicle fusion proteins for their roles during *Drosophila* terminal cell development using RNAi. We uncovered three SNARE proteins, Syntaxin 5 (Syx5), Syntaxin 18 (Syx18), and Syntaxin 1A (Syx1A), that act at distinct steps in the secretory pathway and are required for proper terminal cell morphogenesis. Individual knockdown of each protein resulted in severe reduction of overall branch complexity, phenotypes supported by the disruption of exocyst components and COPII and COPI coat complex assembly proteins. While anterograde (ER-to-Golgi), retrograde (Golgi-to-ER), and Golgi-to-plasma trafficking events all contribute to unicellular branching morphogenesis, the inputs of these portions of the secretory pathway during lumen formation are more complex. Proteins acting at distinct points of ER-to-plasma membrane vesicle delivery had varying contributions to seamless tubulogenesis, whereas Golgi-to-ER retrograde trafficking had minimal inputs during tube formation. Our findings indicate differential inputs on behalf of the secretion pathway during basal and apical membrane growth and maintenance in terminal cells.

## Materials and Methods

### Drosophila Husbandry, Strains, and Crosses

Flies were raised on a standard cornmeal food with sucrose/dextrose (1:2 ratio). Fly crosses were maintained at 25°C unless otherwise stated. The following fly strains were obtained from the Bloomington Stock Center: UAS-*TdTomato* BL# 36328, UAS-*BetaCOP-RNAi* BL#31709, UAS-*Beta’
sCOP-RNAi* BL#31710, UAS-*ZetaCOP-RNAi* BL#28960, UAS-*DeltaCOP-RNAi* BL#31764, UAS-*Syx18-RNAi* BL#26721, UAS-*Sec16-RNAi* V109645, UAS-*Sec31-RNAi* BL#32878, UAS-*2xEGFP* BL#6874, UASp-*RFP-KDEL* BL#30910, UASp-*rfp-golgi* BL#30908, UAS-*Syx1A*-RNAi BL#25811, UAS-*Sec5*-RNAi BL#50556, and UAS-*Sec15*-RNAi BL#27499, *Syx18*^*EYO8095*^/TM3 BL#17430. UAS>*gtaxin* RNAi was kindly provided by the Budnik lab. Expression of Crb was visualized with a GFP-tagged knock-in allele (a generous gift from Hong lab).

To generate pan-tracheal and terminal cell-specific expression, we employed the Gal4-UAS system under either the *breathless* (*btl*) and serum response factor (*srf*) promoters. For branching, lumen, vacuole, Golgi, and ER analysis, srf>gal4, UAS>gfp was used to promote all RNAi knockdown constructs except UAS>Sec31 RNAi and UAS>*gtaxin* RNAi; btl>gal4, UAS>gfp was used for *Sec31* and *Gtaxin*. UASp-rfp-golgi and UAS-rfp-KDEL were crossed into each RNAi background in order to assess Golgi and ER localization. To characterize lum-gfp, btl>gal4, UAS>lum-gfp, UAS>dsRed^NLS^ (2) flies were crossed to the relevant UAS>RNAi fly strain and compared to btl>gal4, UAS>lum-gfp, UAS>dsRed^NLS^, UASp>golgi-rfp (2) control flies. To observe Crbs localization, btl>gal4, crb>crb-gfp, UAS>dsRed (3) flies were crossed to the appropriate UAS>RNAi fly line and compared to btl>gal4, crb>crb-gfp, UAS>dsRed (3) control flies. Control and experimental lum-gfp and crb-gfp crosses were maintained at 18°C in order to avoid early larval lethality that occurs at 25°C. To generate positively marked mutant clones in the tracheal system, UAS-gfpRNAi (Amin’s 2011 screen paper) or MARCM (Lee and Luo 2001) approaches were used. Membrin mutant terminal cells were generated by crossing *Mooncheese*^*1524*^/TM3 TubGal80 flies (Ghabrial Lab, Columbia University) with *ywFLP*^*122*^*;btl-gal4, UAS-gfp, UAS-rfp*^*NLS*^*;gfpiBthstFRT2A*/MKRS flies. FRT82B was recombined onto the Syx18^EYO8095^ chromosome; the resulting recombinant chromosome was then crossed to UAS-gfp, UAS-rfp^NLS^; gfpiBthstFRT82B/MKRS flies. Homozygous mutant *Sec24CD* terminal cells were generated by crossing *Sec24CD*^*KG02906*^ *FRT40A*/CyO (generously provided by Julie Merkle and Gertrud Schüpbach) with *ywFLP*^*122*^*;TubGal80FRT40A*; *btl-gal4* UAS-*gfp* flies.

### Whole Mount Larval Imaging

Wandering third instar larvae were heat-fixed for 5 seconds at 70°C, and mounted in one drop of 60% glycerol before being imaged on either a Leica TCS SP5 X or a Nikon C2 confocal microscope. Images are displayed as either single optical sections or z-stack projections. Image analysis was performed with Fiji. In Figure 4A-D’, ultraviolet light was used to visualize the chitin lining of terminal cells. Chitin autofluorescence has been well documented at emission wavelengths around 450nm (Roshchina 2012).

### Dissection, Fixation, and Immunofluorescence

3rd instar larvae were dissected on a Sylgard pad (ref my 2014 Curr Bio paper) and fixed in 4% paraformaldehyde/1xPBS/0.3% Tween-20-TritonX-100 (1xPBSTwT) for 25 minutes at room temperature. Filets were incubated in primary antibodies overnight and then washed in 1xPBSTwT for 3 x 25 minutes. Samples were incubated in secondary antibodies for 1 hour at room temperature and then washed in 1xPBSTwT for 3 x 25 minutes. Samples were mounted in Aqua-Poly/Mount (PolySciences) before being imaged on a Leica TCS SP5 X (Swarthmore) or Nikon C2 (Princeton) confocal microscope.

The following primary antibodies were used for immunofluorescence: chicken *α*-GFP (1:1000, Invitrogen #A10262), mouse *α*-acetylated tubulin (1:1000, Sigma #T6793), mouse *α*-RFP (1:1000, Abcam #ab65856), and rabbit α-Wkdpep (1:1000, generously provided by Amin Ghabrial). Secondary antibodies utilized in this study include: goat *α*-chicken 488 (1:1000, Life Technologies #A-11039), donkey *α*-mouse 568 (1:1000, Life Technologies #A-10037), donkey *α*-rabbit 647 (1:1000, Life Technologies #A-31573).

### Data Analysis

#### Quantification of terminal cell branching

All branching analysis was performed on whole mount larval images of 4th row (from the posterior) dorsal terminal cells. Three-dimensional traces of each cell were completed using the Simple Neurite Tracer plugin in ImageJ. Sholl analysis was performed to determine branch complexity. The center point was defined as the middle of the cell soma; concentric circles were drawn at 25 µm intervals and the number of branch intersections were calculated for each distance away from the center point. A mixed-effects model with Bonferroni correction was used to determine differences between sample sets.

#### Quantification of lumen, vacuolar, Golgi, and apical protein (Crb) defects

Wild-type and mutant terminal cells were scored for phenotypes via a categorical scoring system. For comparison of phenotypic defects, chi-squared analysis was performed to compare mutant cells to wild-type cells.

## Results and Discussion

### The Secretory Pathway is Required for Proper Branching Morphogenesis in Tracheal Terminal Cells

To examine the membrane trafficking inputs during terminal cell morphogenesis, we performed an RNAi screen against all Qa-SNARE and two Qc-SNARE proteins in *Drosophila melanogaster* using the Gal4-UAS system under the control of the *btl* promoter. Pan-tracheal knockdown for *Syx4,-6,-8,-13,-16,-17* generated viable larvae exhibiting no defects in tracheal terminal cells at the wandering third instar larval stage (data not shown). In contrast, larvae expressing RNAi against *Syx7* displayed terminal cell phenotypes as previously reported (Schottenfeld-Roames, Rosa, and Ghabrial 2014). Furthermore, depletion *of Syx5, Syx18*, and *Syx1A* resulted in 2nd instar larval lethality. To further explore the role of *Syx5, Syx18*, and *Syx1A* during terminal cell formation, we took advantage of the terminal cell-specific gal4 driver, *srf>gal4*. Terminal cell-specific knockdown of all three genes resulted in severely pruned terminal cells (Figure 1A-D). Sholl analysis revealed a statistically significant decrease in branch complexity and outgrowth in *Syx5*-RNAi (*Syx5i*), *Syx18*-RNAi (*Syx18i*) and *Syx1A*-RNAi (*Syx1Ai*) terminal cells (Figure 1E).

**Figure 1.**
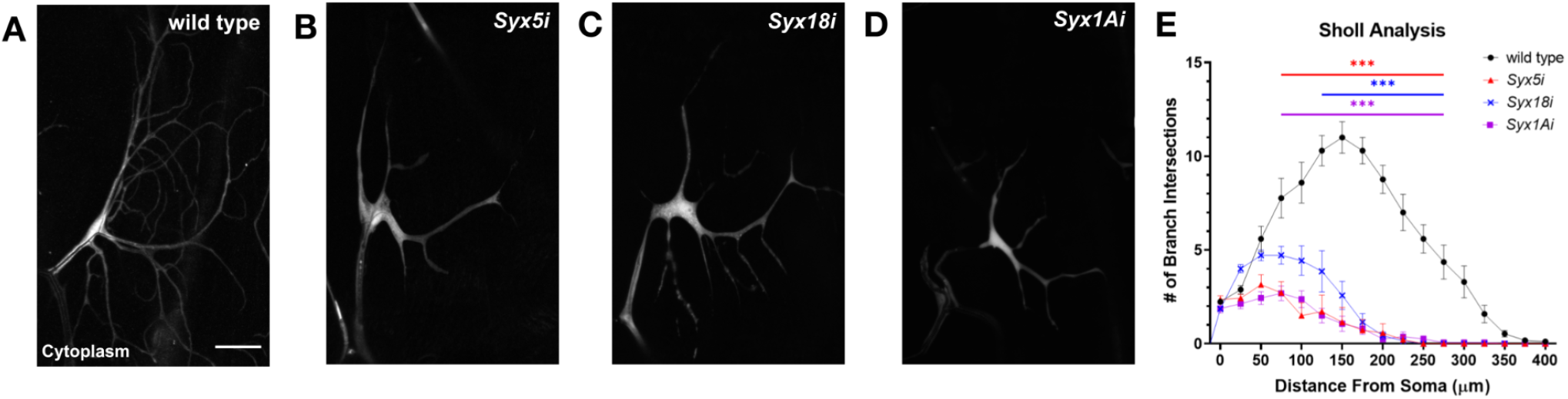
Tracheal terminal cells depleted for Syntaxin 5, 18, and 1A display defects in branching. (A-D) Terminal cells expressing cytoplasmic GFP (white) and a Syntaxin-RNAi. Representative images of wild type (A), *Syx5*-RNAi (B), *Syx18*-RNAi (C), and *Syx1A*-RNAi (D) terminal cells. Scale bar in A represents 50μm. (E) Sholl analysis demonstrates that branching is significantly reduced in 4th dorsal terminal cells *Syx5i, Syx18i*, and *Syx1Ai* dorsal terminal cells between 75-275μm, 125-275μm, and 75-275μm respectively (p<0.001 for all). Symbols and error bars represent mean±SEM. Statistics were calculated using a mixed-effects model followed by Bonferroni multiple comparisons test (wild type, n=17; *Syx5*i, n=14; *Syx18*i, n=7; *Syx1A*i, n=16).

Syx5, Syx18, and Syx1A are Qa-SNAREs that regulate vesicle fusion at distinct points in the secretory pathway (Teng, Wang, and Tang 2001). Syx5 has been shown to localize to ER, Golgi, and COPII-associated vesicles and regulate ER-to-Golgi anterograde trafficking, as well as intra-Golgi transport (Linders et al. 2019). Syx18 is an ER resident SNARE that mediates transport of vesicles from the Golgi to the ER (K. Hatsuzawa et al. 2000), and Syx1A plays a critical role in synaptic vesicle fusion at the plasma membrane (Schulze et al. 1995). As Syx1A is required for fusion of exocytic vesicles with the plasma membrane, it follows that *Syx1Ai* terminal cells display branching phenotypes similar to those shown for exocyst complex mutants (Figure 1D; Jones et al. 2014)). To determine the larger contributions of anterograde and retrograde trafficking to terminal cell morphogenesis, we analyzed terminal cells compromised for the ER exit site protein Sec16 (Watson et al. 2006), the Syx5-interacting Qb-SNARE Membrin (Linders et al. 2019; Ghabrial, Levi, and Krasnow 2011), and COPII and COPI vesicle coat complex proteins. The COPII coat complex promotes anterograde transport of vesicles from the rough endoplasmic reticulum (ER) to the Golgi apparatus, and COPI proteins facilitate the retrograde transport of vesicles budding from the Golgi enroute back to the ER (Bonifacino and Glick 2004). We specifically examined *β-COP, β’-COP, δ-COP*, and *ζ-COP* from COPI complex, and *Sec31, Sec24CD* from COPII complex, as well as other integral proteins, Sec16 and Membrin, for secretion. All terminal cells defective for anterograde and retrograde trafficking generated fewer branches than wild-type control cells (Figure S1). Compromised ER-to-Golgi trafficking led to a range of branching defects (Figure S1 B-D, E), whereas perturbation of COPI-associated components often resulted in terminal cells with a single branch (Figure S1 F-J, K). A notable exception was RNAi against the *δ-COP* subunit of the COPI complex (*δ-COP*i), which presented with a more graded phenotype, exhibiting reduced branching (Figure S1 E) in addition to 32% with a single branch (Figure S1 K). Since further characterization of severely compromised COPI terminal cells was not possible, branched *δ-COP*i cells and *Syx18i* cells were used to assess terminal cell phenotypes in retrograde trafficking defective cells.

### Regulators of both anterograde trafficking and late secretion are required for seamless tubulogenesis

As terminal cells form a subcellular lumen within each branch, we sought to ascertain the role of each portion of the secretory pathway during seamless tubulogenesis. It has been previously shown that different components of the endocytic and lysosomal pathways are critical for seamless tube formation and maintenance (Schottenfeld-Roames, Rosa, and Ghabrial 2014; Schottenfeld-Roames and Ghabrial 2012; Nikolova and Metzstein 2015; Francis and Ghabrial 2015). Other studies have suggested a role for the exocyst complex during this process (Jones and Metzstein 2011). To assess the inputs of the secretory pathway during seamless tube formation in terminal cells, we utilized the α-Wkd apical membrane marker in representative sets of knockdowns (Figure 2) of anterograde (red), retrograde (blue), and late-stage (purple) secretory pathway components (Schottenfeld-Roames and Ghabrial 2012). Cells compromised for Golgi-plasma membrane transport and fusion displayed a myriad of lumenal phenotypes, including: multiple lumens within one branch (Figure 2B-B”), truncated lumens that do not extend out to branch tips (Figure 2C-C”), discontinuous lumens (Figure 2D-D”), and disorganized lumens consisting of tangles or abnormal membrane morphology (Figure 2E-E’’’’). In contrast to these dramatic phenotypes, knockdown of regulators for ER-to-Golgi and Golgi-to-ER trafficking showed diminished effects on tube morphogenesis (Figure 2A-E’’’’). Notably, only knockdown of Syx5, a SNARE protein required for the final step of anterograde trafficking and subsequent intra-Golgi vesicle transport, resulted in phenotypes mirroring those of compromised late-stage secretion (Figure 2A-E”’’). Unexpectedly, terminal cells lacking the COPII coat protein Sec31 exhibited a high incidence of microdilations at the lumenal membrane reminiscent of the lumenal cysts observed in endocytic mutants (Figure 2F-F”; (Schottenfeld-Roames, Rosa, and Ghabrial 2014); this phenotype was also seen to a lesser extent in *Sec16* depleted cells. This is the first demonstration that members of the secretory pathway contribute to subcellular tube architecture. These results may imply crosstalk between secretory and endocytic pathways. Indeed, previous studies have implicated direct ER involvement in phagocytosis, a process which internalizes extracellular material similar to endocytosis (Kiyotaka Hatsuzawa et al. 2006). We conclude that all secretory pathway components do not have equal inputs into seamless tubulogenesis in tracheal terminal cells. Analysis of apical membrane trafficking shown here has demonstrated variable, protein-specific inputs into tube formation for regulators of ER-Golgi-plasma membrane vesicle transport, with less weight assigned to retrograde trafficking during this process.

**Figure 2.**
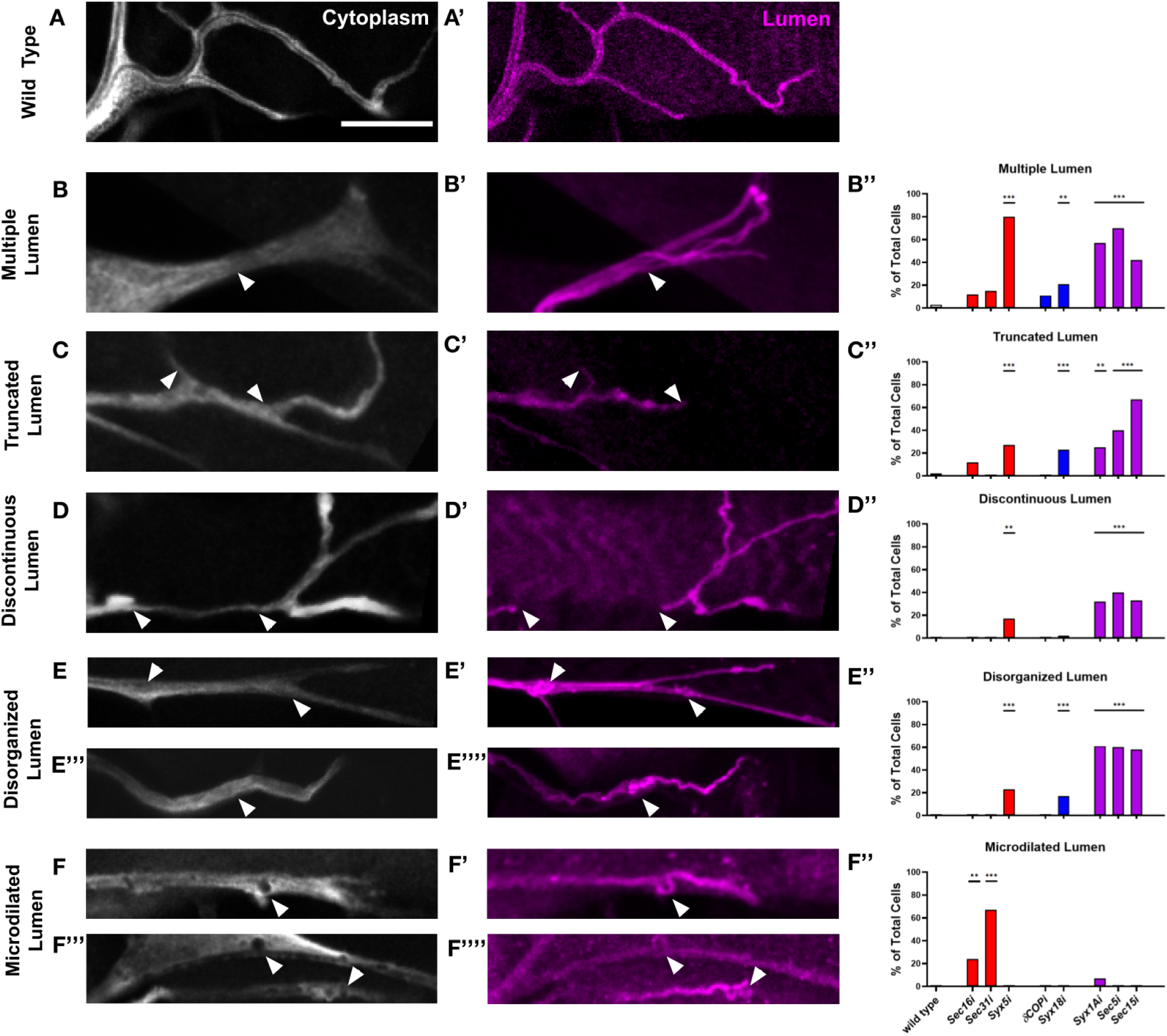
Secretory pathway components are differentially required for lumen morphology, growth, and maintenance in terminal cells. (A-F’’) Representative images of fixed 3rd instar larvae immunostained for cytoplasmic GFP (white) and the lumenal membrane (α-Wkdpep, magenta). Unlike the single, smooth lumen that transverses each wild-type branch (A-A’), cells compromised for anterograde, retrograde, and late secretion result in various lumen phenotypes, including multiple lumens within a single branch (arrowheads in B-B’) truncated lumens that do not extend to the tip of the branch (arrowheads in C-C’), discontinuous lumens (arrowheads in D-D’), disorganized tangles or clumps of lumen (arrowheads in E-E’’’’), and lumens that exhibit membrane microdilations (arrowheads in F-F’). Scale bar in A represents 20μm. (B’’-F’’) Terminal cells depleted for the secretory pathway proteins indicated on the x axis were scored for each of the five aberrant lumen phenotypes. Bar color represents anterograde (red), retrograde (blue), and late (purple) secretory pathway affected cells. Chi-square analysis for individual knockdowns was performed comparing wild-type to RNAi cells (**: p<0.01; ***: p<0.001, wild type, n= 61; *Sec16i*, n= 17; *Sec31i*, n= 27; *Syx5i*, n= 30; *δCOPi*, n= 19; *Syx18i*, n=66; *Syx1Ai*, n=28; *Sec5i*, n=10; *Sec15i*, n=12.)

### Knockdown of anterograde and retrograde trafficking results in vacuole-like structures and aberrant localization of secretory organelles in terminal cells

Although we observed varying effects on tube formation when anterograde and retrograde transport were disrupted, cytoplasmic architecture was consistently affected in these cells. The cytoplasm of wild-type terminal cells is largely uniform when labelled with a cytoplasmic fluorescent reporter protein (Figure 3A). Occasionally, small “bubbles” of cytoplasmic exclusion are observed adjacent to branch points or within a branch extension (Figure 3B). These spheres have been shown to fill with a secreted form of GFP, suggesting they are intermediates of the secretory pathway (Ghabrial, Levi, and Krasnow 2011). When regulators of anterograde and retrograde trafficking were disrupted, these cytoplasm-excluding spheres became more prevalent and pronounced. Affected cells displayed a range of vacuole-like structures from numerous small cytoplasm-excluding bodies (Figure 3C) to large vacuoles in the cell soma, similar to those observed in Membrin mutants (Figure 3D; (Ghabrial, Levi, and Krasnow 2011). In *Sec31i* terminal cells, a large fraction of these structures appeared contiguous with the lumen (Figure 3E), consistent with the earlier observation that *Sec31* depletion results in apical membrane microdilations (Figure 2G-G’’). Knockdowns of *Sec16, Sec31, Syx5*, and *Syx18* all resulted in a significantly greater proportion of cells containing vacuole-like structures compared to controls (Figure 3F). Terminal cells mutant for *Syx18*, or ones that expressed an alternative RNAi against *Syx18*, confirmed the vacuolar phenotype (Figure S2). Unlike terminal cells compromised for ER-Golgi transport, large, proximal vacuoles were not observed in *Syx1Ai, Sec5i*, or *Sec15i* cells (data not shown); *Syx1Ai* and *Sec15i* cells did exhibit a mild increase in small cytoplasmic-excluding bodies compared to wild-type (Figure 3F).

**Figure 3.**
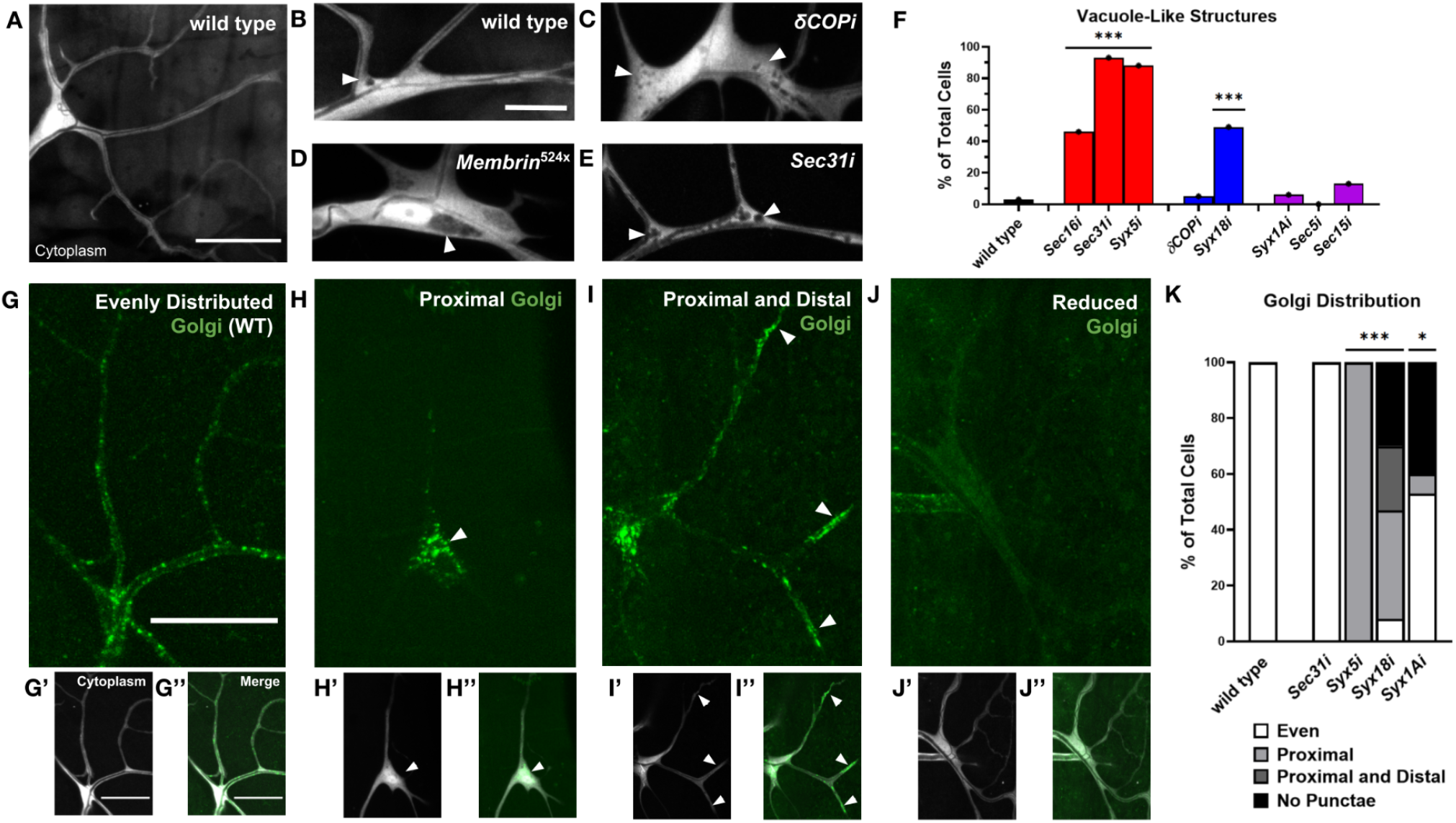
Perturbation of the secretory pathway leads to the accumulation of intracellular vacuoles and disruptions to Golgi architecture in terminal cells. (A-E) Terminal cells expressing cytoplasmic GFP (white). Wild type cells (A) have continuous cytoplasm with occasional small vacuole-like structures at branch points (arrowhead in B). Terminal cells compromised for secretion present with large vacuoles in the cell soma (arrowhead in D) and/or an increased number of small “bubbles” of cytoplasmic exclusion throughout the terminal cell (arrowheads in C and E). Representative images of vacuole phenotypes are shown for *δ-COP* RNAi (C) and *Membrin*^524^ (D) with percentages of wild type and RNAi knockdowns displaying these phenotypes illustrated in F. Cytoplasmic exclusions present in *Sec31*-RNAi terminal cells are frequently in close proximity to the subcellular lumen (E). (F) Frequency of cytoplasmic exclusions in terminal cells is quantified and stratified by early anterograde (red), early retrograde (blue) and late (purple) secretion pathway knockdowns. Chi-square analysis for individual mutants was performed comparing wild-type to mutant cells (***: p<0.001, wild type, n= 64; *Sec16i*, n= 67; *Sec31i*, n= 41; *Syx5i*, n=17; *δCOPi*, n= 92; *Syx18i*, n=51; *Syx1Ai*, n=33; *Sec5i*, n=5; *Sec15i*, n=8. (G-J’’) Terminal cells expressing cytoplasmic GFP (white) and GalT-RFP (Golgi, green). Wild-type terminal cells (G-G’’) have evenly distributed Golgi punctae throughout the terminal cell. *Syx18*-RNAi terminal cells (H-J”) do not exhibit a wild type Golgi distribution. *Syx5, Syx18, and Syx1A* knockdown cells display a variety of phenotypes ranging from enrichment of Golgi in the cell soma (defined here as proximal, arrowheads in H-H’’) with an absence of punctae in the branch extensions, aggregates of Golgi in both proximal and distal regions (branch tips, arrowheads in I-I’’), and a global reduction of Golgi punctae (J-J’’). (K) Golgi phenotypes were quantified across all secretory pathway knockdowns with chi-square analysis comparing WT to mutant cells (*: p<0.05; ***: p<0.001, wild type, n=30; *Sec31i*, n=22; *Syx5i*, n=17; *Syx18i*, n=53; *Syx1Ai*, n=15). *Sec31*-RNAi cells exhibit a wild type Golgi organization. Scale bars in A represent 50μm, B-E represent 20μm, and G-J represent 50μm.

We reasoned that in mutants where vesicle trafficking between the ER and Golgi is blocked, these cytoplasm-excluding compartments might be a consequence of disorganized ER-Golgi compartments. It has been shown that in mammalian cells, knockdown of *Syx5* and *Syx18* results in changes to Golgi and ER organization, respectively (Suga et al. 2005; Iinuma et al. 2009). To determine whether Golgi architecture was disrupted in our protein knockdowns, we utilized the Gal4-UAS system to express an RFP-tagged version of the trans-Golgi marker GalT under either the *btl* (pan-tracheal) or the *srf* (terminal-cell specific) gal4 driver. While all wild-type terminal cells expressed evenly distributed fluorescent Golgi punctae throughout the cell body and branches (Figure 3G,K), RNAi knockdowns of secretory pathway components resulted in several phenotypes, including proximally-restricted GalT punctae (Figure 3H), GalT localization proximally and at distal tips (Figure 3I), or reduced GalT punctae (Figure 3J). 100% of *Syx5i* cells displayed large aggregates of GalT in the cell soma (Figure 3K) and unlike other mutants, some of these aggregates did overlap with pockets of cytoplasmic exclusion. *Syx18i* terminal cells showed a more variable effect on Golgi distribution; a combined 62% displayed either proximally-restricted Golgi punctae or enrichment of GalT-positive punctae in the cell soma and branch tips, with 30% of cells showing reduced Golgi puncta throughout the cell (Figure 3K). Consistent with a small increase in cytoplasmic “bubbles” in *Syx1Ai* terminal cells, 7% of *Syx1Ai* cells displayed cell soma restricted GalT:RFP and 53% of *Syx1Ai* cells displayed normal Golgi organization. Interestingly, the other 40% of cells had a reduction in Golgi punctae throughout the cell (Figure 3G,J,K).

Utilizing a fluorescent ER retention signal peptide (KDEL) we observed a similar disruption in ER organization in *Syx18i* terminal cells, with 32% displaying cell soma enrichment, 14% showing both distal and proximal localization, 30% displaying a diffuse signal lacking punctae, and only 36% with an even ER localization (Figure S3A-E). In *Syx1Ai* terminal cells, 25% of cells displayed cell soma restricted KDEL; 44% of cells displayed normal KDEL organization, and 31% had diffuse KDEL throughout the cell (Figure S3E). Since Syx1A is thought to act downstream of both the ER and Golgi to promote vesicle fusion at the plasma membrane, it is surprising that both ER and Golgi architecture were affected in *Syx1Ai* terminal cells.

Golgi organization was not altered in *Sec31i* terminal cells, providing additional support that the lumen-adjacent vacuoles observed in these cells were primarily dilations of the lumenal membrane and not indicative of gross abnormal ER-Golgi associated membrane-bound compartments (Figure 3). Taken together, these data suggest that when secretion is compromised, ER and Golgi aggregate in the cell soma and branches, resulting in abnormal organelle localization that is consistent with the vacuolar phenotypes we have observed.

Cell types with unconventional cell shapes, such as muscles, oligodendrocytes and neurons, have specialized golgi structures termed golgi outposts, that support the development of their unique forms (Valenzuela et al. 2020). In mammalian and *Drosophila* da neurons, golgi outposts reside at dendritic branch points and shafts and serve as acentrosomal microtubule-organizing centers (MTOCs) (Ye et al. 2007; Zhou et al. 2014). These satellite golgi structures establish microtubule polarity within dendritic branches and facilitate dendrite growth through local secretion and protein modification; whether golgi outposts act in membrane addition at these sites is still unclear (Valenzuela et al. 2020). In wild type tracheal terminal cells, we have not observed Golgi enrichments at terminal cell branch points, but rather even distribution of t ER and GalT-marked Golgi punctae in the cell soma and branches. Since our data strongly suggests that this spatial organization is critical for basal and apical membrane expansion, the question remains whether these uniformly-spaced Golgi vesicles are structurally and functionally equivalent to the Golgi outposts observed in oligodendrocytes and da neurons. It is possible that outposts do exist in Drosophila terminal cells, but that the GalT trans-Golgi marker may be insufficient to identify the relevant subset among these multicompartmental structures (Zhou et al. 2014). *Cis*, medial, and additional *trans*-Golgi markers will need to be screened to assess whether Golgi outposts exist in the same form and function in terminal cells as in these other distinctively shaped cells. It is intriguing to postulate that discrete Golgi compartments act as additional sites for microtubule nucleation in terminal cells. These presumptive MTOCs could be instructive for branching, in contrast to the apically nucleated subset of microtubules that drive tube extension (Gervais and Casanova 2010; Schottenfeld-Roames and Ghabrial 2012).

It remains unclear as to why ER and Golgi aggregate in proximal and distal regions of terminal cells when secretion is disrupted. One possibility is that microtubule nucleation is abrogated in these mutant backgrounds, leading to aberrant organization of secretory organelles. Because membrane trafficking is a dynamic process, it will be especially informative to follow movement of the ER and Golgi compartments during terminal cell morphogenesis in real-time in wild type and *Syx* mutant terminal cells. Live-imaging has proved easier in embryonic terminal cells, where multivesicular bodies have recently been shown to direct local transcytosis of both apical and basal components at the cell tip (Mathew et al. 2020). Since the behavior of secretory organelles has not yet been addressed during membrane expansion in terminal cells, these studies could provide new insights into whether ER and Golgi organelles have direct interactions with the apical and basal domains during branch and lumen maturation.

### Secretion of Lumenal and Apical Membrane-bound Proteins is Compromised upon depletion of Syx5, Syx18, and Syx1A

To determine whether secretion at the lumenal membrane is disrupted on depletion of *Syx5i, Syx18i*, and *Syx1Ai* in terminal cells, we examined a secreted reporter protein, Lum-GFP (Ghabrial, Levi, and Krasnow *2011*), in wild-type and knockdown. Because the lumens of wild-type terminal cells are air-filled, Lum-GFP does not accumulate inside terminal cell tubes; thus, most Lum-GFPfluorescence is observed as a thin line of signal flanking the lumen (Figure 4A-A’’’, (Schottenfeld-Roames, Rosa, and Ghabrial 2014)). In striking contrast, *Syx5i* and *Syx18i* cells displayed ectopic accumulation of Lum-GFP in the cell soma (Figure 4B-C’’’), suggesting that early secretory pathway intermediates build up inside the terminal cells. Importantly, we did not observe these cell soma aggregates in *Syx1Ai* terminal cells, (Figure 4D-D’’’). All three RNAi knockdowns presented varying degrees of air-filling defects, visualized by the absence of the dark line normally illuminated by brightfield microscopy (Figure 4A’’’-D’’’). In all such cases, the lumenal chitin matrix was visible by UV light (magenta), indicating that lumen was present but was fluid-filled rather than air-filled. When secretion is intact, lum-GFP brightly fluoresces in lumens of fluid-filled tubes (Ghabrial, Levi, and Krasnow 2011). Lum-GFP did not highlight the chitin-marked lumens in *Syx18i* cells (Figure 4C-C’’’) and displayed patchy fluorescence in the tubes of *Syx5i* and *Syx1Ai* cells (data not shown and Figure 4D’’). These results indicate that lumenal components were not being secreted properly in *Syx5i, Syx18i*, and *Syx1Ai* cells, and that this block in secretion occurred at different steps in the pathway.

**Figure 4.**
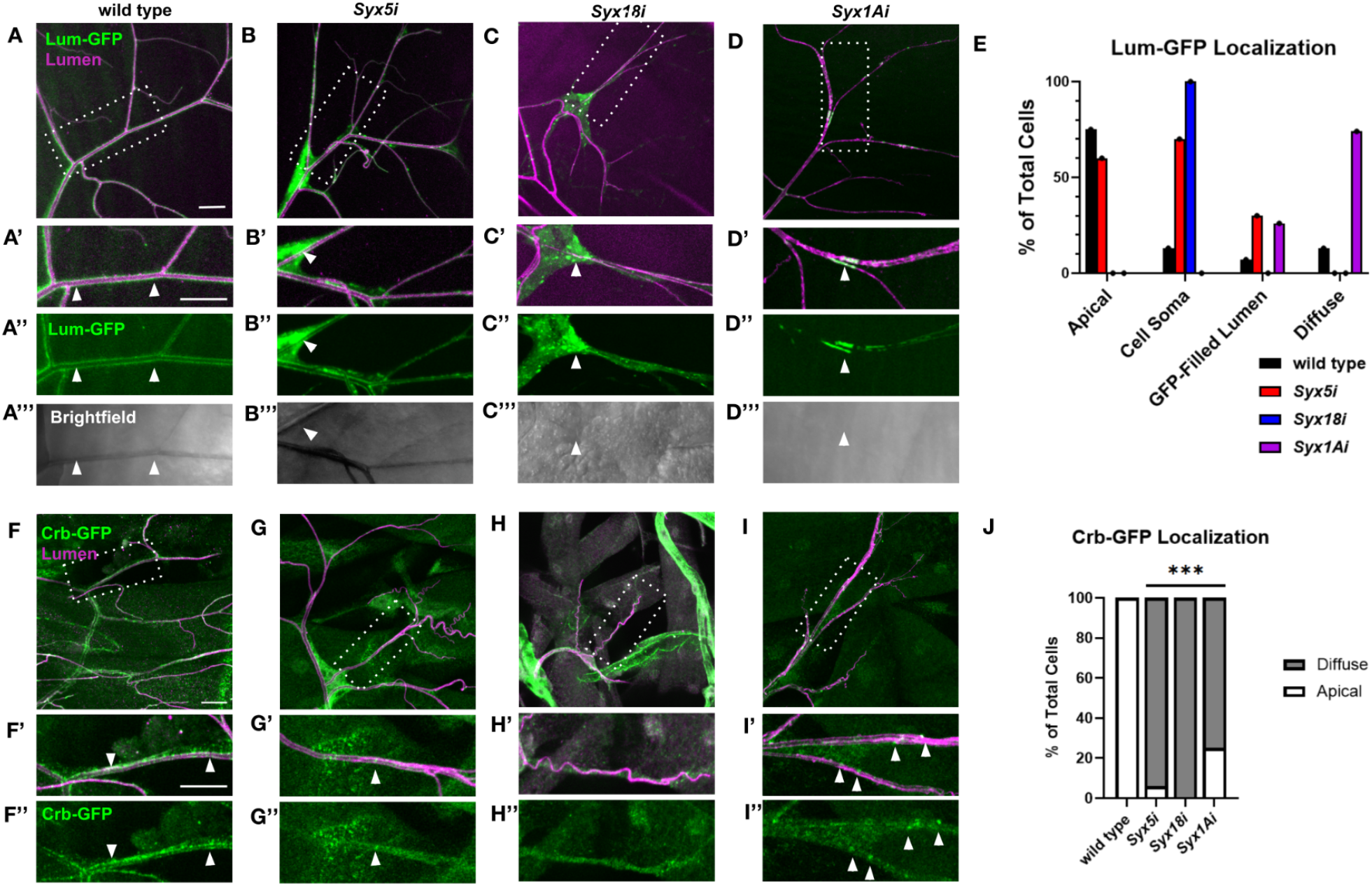
Secretion of lumenal and apical membrane proteins is compromised in *Syx5i, Syx18i, and Syx1Ai* terminal cells. (A-D’’’) Composite images (A-D) of terminal cells expressing a secreted form of GFP (Lum-GFP, green) and the apical chitin matrix (lumen, magenta). White boxes in A-D are magnified in A’-D’. Lum-GFP alone is shown in A’’-D’’ and the gas-filled lumen (grey) is visible in A’’’-D’’’. Lum-GFP flanks the lumen of gas-filled tubes in wild type cells (A, arrowheads in A’-A’’’) but accumulates in the cell soma of *Syx5*i (B, arrowheads in B’-B’’’) and *Syx18*i (C, arrowheads in C’-C’’’) cells. Syx18i cells do not gas-fill properly; however, Lum-GFP is not secreted into the fluid-filled tube. *Syx1A*i terminal cells do not accumulate Lum-GFP in the cell soma; instead, they show faint, diffuse staining of GFP throughout the cell body (D-D’’’). Isolated patches of Lum-GFP are present in *Syx1A*i fluid-filled tubes (arrowheads, D’-D’’’). (E) Quantification of Lum-GFP localization in discrete genotypes: wild-type (black, n=17), *Syx5*i (red, n=10), *Syx18*i (blue, n=15), and *Syx1A*i (purple, n=19). Phenotypes are not mutually exclusive. (F-I’’’) Terminal cells immunostained for Crb:GFP (green) and α-Wkdpep (lumen, magenta). Composite images are shown in F-I; white boxes are magnified in F’-I’. Discrete foci of Crbs:GFP decorate the apical membrane of terminal cells (F’,F’’). In contrast, Crbs:GFP is predominantly cytoplasmic in *Syx5*i (G-G’’’), *Syx18*i (H-H’’), and *Syx1A*i (I-I’’) terminal cells. Arrowheads in F’,F’’,I’,I’’ highlight Crbs punctae at the apical membrane. (J) Quantification of Crbs-GFP apical localization with chi-square analysis comparing WT to mutant cells (***: p<0.001, wild type, n=7; *Syx5i*, n=16; *Syx18i*, n=10; *Syx1Ai*, n=12). Scale bars in A,A’,F,F’ represent 20μm.

To test whether these secretion defects also affected the targeting of lumenal membrane proteins, we examined the expression of the apical polarity protein Crumbs. Crumbs is a member of the conserved Crumbs (Crbs) / Stardust (Sdt) / PALS-1 Associated Tight Junction Protein (dPatj) apical polarity complex and has been shown to have varying inputs during terminal cell morphogenesis (Schottenfeld-Roames, Rosa, and Ghabrial 2014; JayaNandanan, Mathew, and Leptin 2014; Jewett and Prekeris 2018). In wild-type terminal cells, Crb-GFP localizes along the apical membrane exhibiting a mostly punctate appearance (Figure 4F-F’’, (Schottenfeld-Roames and Ghabrial 2012)). In *Syx5i, Syx18i* and *Syx1Ai* cells, Crb-GFP localization was significantly altered from wild-type cells (Figure 4J). In these secretion-compromised backgrounds, Crb was primarily diffuse in the cell cytoplasm (Figure 4G-I’’). As Syx1A regulates the last step in the exocytic process, it is not surprising that approximately 30% of *Syx1Ai* cells efficiently localized Crb at the apical membrane (Figure 4I-I’’, J). From these data, we conclude that a subset of lumenal membrane proteins do not localize properly when anterograde, retrograde, and late-stage secretion is disrupted. Nonetheless, lumen and chitin deposition still occur despite aberrant secretion, potentially indicating involvement of compensatory trafficking components and pathways for specific cargo.

Terminal cell morphogenesis and migration relies on the targeted secretion of a plethora of transmembrane and extracellular matrix proteins. The delivery of these proteins must be exquisitely coordinated, as both the apical and basal plasma membranes are simultaneously growing and extending toward hypoxic cues. For example, the apical cuticle lining of these tracheal tubes is dependent on the continuous delivery of components required for chitin fibrillar formation (Rosa, Metzstein, and Ghabrial 2018). In epidermal cells, Syx1A is not required for the apical secretion of the chitin modifying proteins Serpentine, Piopio, and Knk, however, outer cuticle organization is nonetheless altered in *Syx1A* mutant larvae (Moussian et al. 2007). Similarly, chitin fibrils still line the abnormal lumens of larval *Syx1Ai* terminal cells, arguing for a role for Syx1A in the selective secretion of extracellular matrix molecules in both epidermal and tracheal cell types. These data suggest that other Q-SNARE proteins must act in parallel to Syx1A at the apical plasma membrane to ensure effective cuticle differentiation. This is likely true for many, if not all, the SNARE proteins analyzed in this study, and may explain why *Syx18i* and *δ-COPi* cells do not show defects in lumen formation when secretion at the apical membrane is at least partially compromised (*δ-COPi* data not shown). Indeed, a mechanism for differential secretion of structural components to the apical membrane has already been documented in *Drosophila* tracheal cells (JayaNandanan, Mathew, and Leptin 2014). Future studies will need to be directed at specialized ER-to-Golgi regulators, such as Sec22 and Tango-1, which have been implicated in the selective secretion of basal ECM proteins, to obtain a more complete picture of the coordinated efforts amongst secretion factors to deliver the myriad of materials required for seamless apical invagination (Linders et al. 2019; Lerner et al. 2013). Depletion of Tango-1 has been shown to cause defects in terminal cell shape, due at least in part to the activation of the ER stress response (Ríos-Barrera et al. 2017).

Our data also confirms previous demonstrations that the targeting of key apical polarity proteins to the lumenal membrane of larval terminal cells is dependent on the exocyst complex (Jones et al. 2014). Interestingly, the absence of lumen phenotypes in *Syx18i* cells, which lack apically localized Crb protein, calls into question the necessity of Crb during subcellular tube formation. These results lend support to our previous findings that Crb is not required for terminal cell morphogenesis but can cause catastrophic consequences to seamless tube morphology when present in excess at the lumenal membrane (Schottenfeld-Roames, Rosa, and Ghabrial 2014). The secreted cargo instructing basal membrane extension remains even more elusive. It is known that developing branch extensions rely heavily on integrin linker proteins to maintain attachment to the basal membrane and sustain robust connections with the internal lumen (Levi, Ghabrial, and Krasnow 2006). Loss of integrins at basal branch extensions is likely one contributing factor for the pruned phenotype in secretion-compromised terminal cells, but determining the full complement of basal-destined proteins supporting unicellular branching morphogenesis is an open area for discovery.

## Conclusion

Here we examined the contributions of discrete steps of the secretion pathway during terminal cell morphogenesis. When components from anterograde, retrograde, or late-stage secretion are compromised, unicellular branching morphogenesis is robustly perturbed. Large quantities of membrane, along with a diverse set of membrane fusion molecules, secreted extracellular matrix (ECM) and ECM-interacting proteins are surely required to achieve the large surface area attained by terminal cells. With this in mind, our data suggest that the secretion pathway takes an “all hands on deck” approach to supplying the cell with the membrane and cargo that it requires for such a significant cell shape transformation. Conversely, these pathways do not contribute equally when building and shaping the cell’s intracellular lumen. Retrograde trafficking components have minimal inputs into seamless tube formation; the ER-Golgi transport regulators, Sec16 and Sec31, control tube size and shape, similar to proteins in the endocytic pathway, whereas the Golgi Syx5 SNARE, the plasma membrane SNARE Syx1A, and exocyst complex subunit Sec15 promote tube growth and maintenance. Our data suggest that elucidating the complex mechanism of apical invagination and/or cell hollowing will require a comprehensive interrogation of factors regulating the flow of protein transport from the ER to the nascent apical membrane.

## Acknowledgements

The authors would like to thank Jessica Sullivan-Brown and Steve DiNardo for comments on the manuscript, as well as the Swarthmore Biology Department for providing an interactive, scholarly atmosphere in which to conduct these studies. We also thank the 2018 Spring Core Lab students at Princeton University for thoughtful discussions about *Syx1A* and *Syx5* RNAi knockdown phenotypes.

---

**Supplemental Figure 1.**
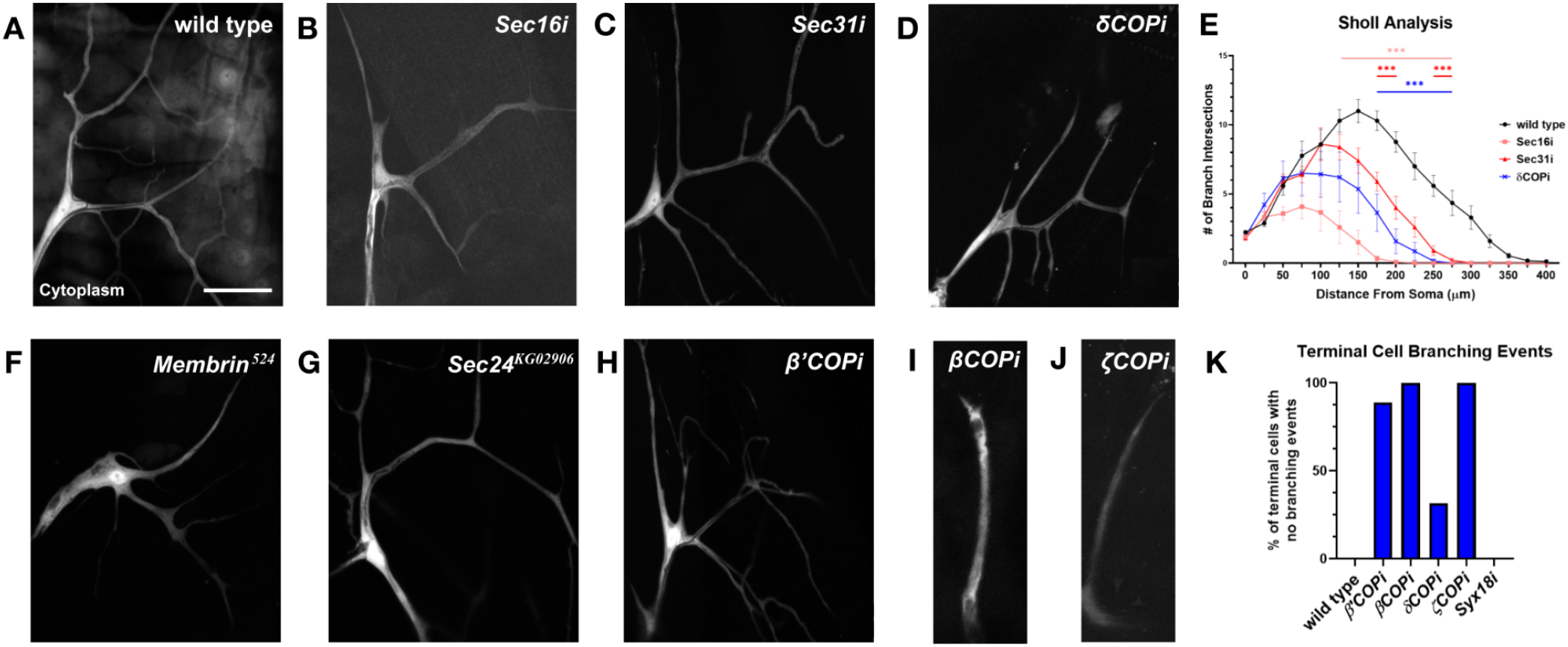
COPII, COPI, and ER exit site proteins are required for proper terminal cell branching. (A-D, F-J) 4th row dorsal terminal cells expressing cytoplasmic GFP (white). *Sec16*-RNAi (B), *Sec31*-RNAi (C), and *δ-COP*-RNAi (D) depleted cells have reduced branching compared to wild type (A). Scale bar in A represents 50 μm. (E) Sholl analysis illustrates that branching is significantly reduced in *Sec16-RNAi, Sec31-RNAi*, and *δ-COPi* cells between 125-275μm, 175-275μm, and *1*75-275μm respectively (p<0.001 for all). Symbols and error bars represent mean±SEM. Statistics were calculated using a mixed-effects model followed by Bonferroni multiple comparisons test (wild type, n=17; *Sec16*-RNAi, n= 12; *Sec31*-RNAi, n= 10; *δ-COP*-RNAi, n= 14). *Membrin*^524^ (F) *Sec24*^*KG02976*^ (G), and *β’-COP*-RNAi (H) terminal cells also display branching defects. RNAi knockdown of COPI subunits, such as shown for *β-COP*-RNAi (I), and *ζ-COP*-RNAi (J) resulted in cells with no branching events. (K) Prevalence of COPI-RNAi terminal cells that did not branch. The retrograde trafficking regulator, Syx18-RNAi, is included for comparison purposes. Wild type, n= 64; *β’-COP*i, n= 18; *β-COP*i, n= 29; *δ-COP*i, n= 92; *ζ-COP*i, n= 12; *syx18*i, n= 51.

**Supplemental Figure 2.**
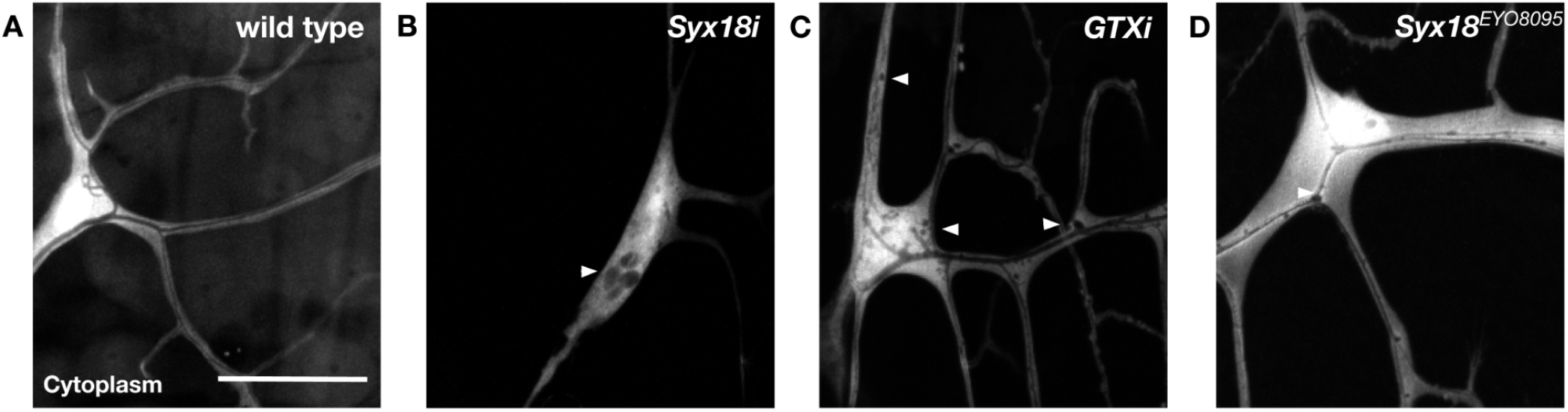
Reduced Syx18 activity results in intracellular vacuole formation in terminal cells. (A-D) Terminal cells expressing cytoplasmic GFP (white). The cytoplasm of wild type terminal cells is largely vacuole free (A). *Syx18*-RNAi expressing terminal cells are severely pruned and display large regions of cytoplasmic exclusion in the cell soma (arrowhead, B). Cells expressing *Gtaxin* (GTX) RNAi (C), an independent RNAi line against *Syx18*, and homozygous *Syx18*^EYO8095^ (D) mutant cells have milder branching phenotypes and small cytoplasm-excluding structures throughout the cell (arrowheads in B and C). Scale bar in A represents 50μm.

**Supplemental Figure 3.**
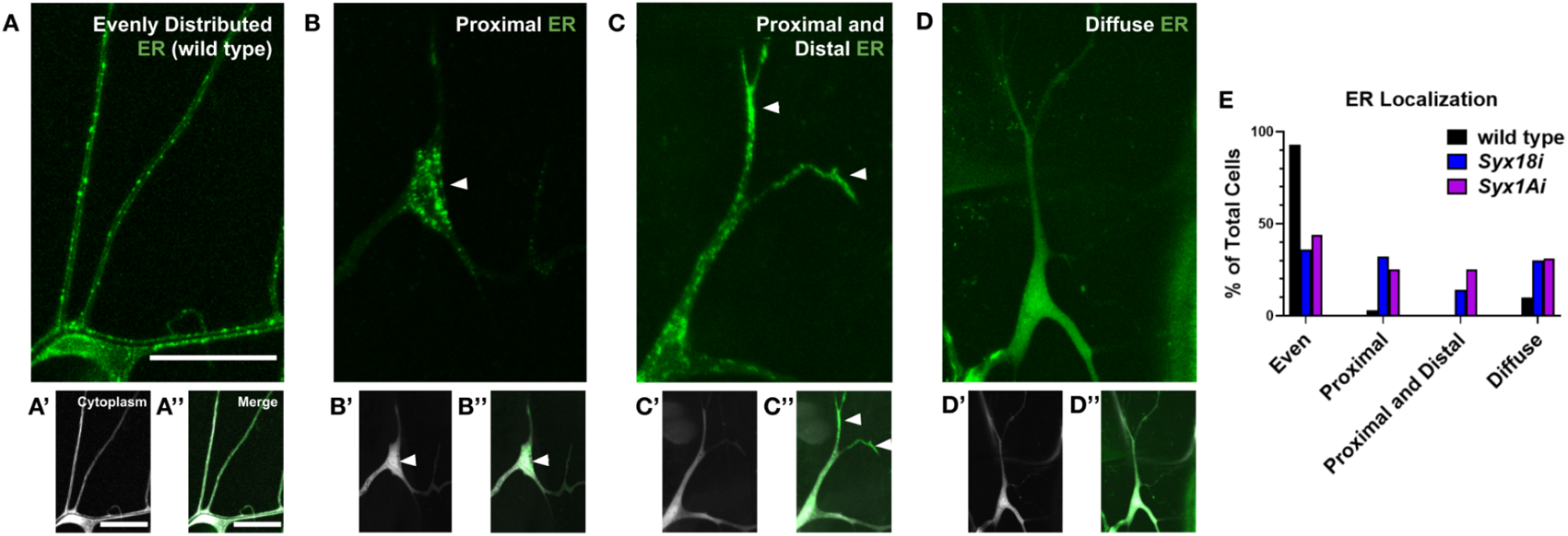
The spatial organization of the endoplasmic reticulum (ER) is altered in *Syx18i* and *Syx1Ai* terminal cells. **(A-D’’)** Terminal cells expressing the ER reporter, KDEL-RFP (green), and cytoplasmic GFP (white). Wild-type terminal cells (A) display evenly distributed ER puncta throughout the body of a terminal cell. *Syx18*-RNAi (B-D’’,E) and *Syx1A*-RNAi cells (E) exhibit an array of phenotypes with respect to ER organization, including enrichment of ER punctae in the cell soma at the expense of branches (arrowheads in B-B’’), ER enrichment in both the cell soma and distal branch tips (arrowheads in C-C’’), and diffuse reporter expression with few defined punctae (D-D’’). (E) ER phenotypes were quantified across all secretory pathway knockdowns. Wild type, n= 30; *Syx18i*, n= 44; *Syx1Ai*, n= 66. Scale bars in A-A’’ represent 50μm.

